# Deep sncRNA-seq of the PPMI cohort to study Parkinson’s disease progression

**DOI:** 10.1101/2020.06.01.127092

**Authors:** Fabian Kern, Tobias Fehlmann, Ivo Violich, Eric Alsop, Elizabeth Hutchins, Mustafa Kahraman, Nadja Liddy Grammes, Pedro Guimarães, Christina Backes, Kathleen Poston, Bradford Casey, Rudi Balling, Lars Geffers, Rejko Krüger, Douglas Galasko, Brit Mollenhauer, Eckart Meese, Tony Wyss-Coray, David Wesley Craig, Kendall Van Keuren-Jensen, Andreas Keller

## Abstract

Coding and non-coding RNAs have diagnostic and prognostic importance in Parkinson’s diseases (PD). We studied circulating small non-coding RNAs (sncRNAs) in 7, 003 samples from two longitudinal PD cohorts (Parkinson’s Progression Marker Initiative (PPMI) and Luxembourg Parkinson’s Study (NCER-PD)) and modelled their influence on the transcriptome. First, we sequenced sncRNAs in 5, 450 blood samples of 1, 614 individuals in PPMI. The majority of 323 billion reads (59 million reads per sample) mapped to miRNAs. Other covered RNA classes include piRNAs, rRNAs, snoRNAs, tRNAs, scaRNAs, and snRNAs. De-regulated miRNAs were associated with the disease and disease progression and occur in two distinct waves in the third and seventh decade of live. Originating mostly from a characteristic set of immune cells they resemble a systemic inflammation response and mitochondrial dysfunction, two hallmarks of PD. By profiling 1, 553 samples from 1, 024 individuals in the NCER-PD cohort using an independent technology, we validate relevant findings from the sequencing study. Finally, network analysis of sncRNAs and transcriptome sequencing of the original cohort identified regulatory modules emerging in progressing PD patients.

## Introduction

Parkinson’s disease (PD) is the second most common neurodegenerative disease worldwide^1^ but the exact causes for the disease and its progression remain largely unknown^2^. PD appears to result from a complicated interplay of genetic and environmental factors, which are affecting fundamental cellular and molecular processes^3^. The diagnosis by means of the often very heterogeneous symptoms is based on clinical criteria^4^, making validated diagnostic and prognostic biomarkers an unmet demand in order to support novel therapeutic developments. The rapid development of high-throughput screening techniques and declining experimental costs fostered larger PD studies such as Genome- or Transcriptome Wide Association Studies to discover potential marker genes^5–9^. On the search for low-invasive biomarkers, transcriptome analyses based on a variety of bio-fluids such as blood^10–12^ or cerebrospinal fluid^13^ have been conducted. Until recently, these studies were primarily focused on the coding transcriptome, obtruding the question about the role of non-coding RNAs in PD^14^.

As part of the sncRNAs, microRNAs play a remarkable role. Due to their stability, diagnostic and prognostic information content, and the well-understood downstream targeting effects on gene expression they are promising candidates, also in the context of PD^15, 16^. Already in 2011, we provided evidence that blood-borne miRNAs can serve as specific markers for human pathologies^17^. Based on the salient cross-disease results we focused on neurodegenerative disorders such as Alzheimer’s disease^18–20^ and malignant tumor diseases such as lung cancer. In the latter case, we performed one of the largest diagnostic miRNA studies to date, covering over 3, 000 patients and controls^21^.

Advanced biomarker studies however require carefully designed cohorts. Already a few large-scale PD studies aiming to advance diagnosis, prognosis and therapeutics fulfil respective requirements^22–24^. Among them, the Parkinson’s Progression Marker Initiative (PPMI) is a multi-cohort, longitudinal observational study designed to discover and validate objective biomarkers of Parkinson’s^25^. The PPMI project constitutes a global effort of 33 clinical sites in 11 countries with regular study participant assessments (Figure 1a). It also features comprehensive clinical phenotyping to observe hundreds of characteristics of the known subtypes of the disease, such as the idiopathic and genetic forms. Further, longitudinal biosampling following rigorous Standard Operating Procedures (SOPs) is performed to set a framework for the discovery and validation of early-onset and prognostic biomarkers. To identify potential non-coding RNA and transcriptomic markers in PPMI we performed RNA-seq on blood samples drawn at each clinical visit. For short and long RNAs, we carried out optimized assays and sequenced separate aliquots from the same blood samples for paired RNA analyses. Here, we present the evaluation of the sncRNA-seq fraction for disease detection and progression tracking. We examine the potential of different classes of small RNAs but emphasise the role of miRNAs. We also validate relevant findings for miRNAs on the Luxembourg Parkinson’s Study in the framework of the National Centre for Excellence in Research on PD (NCER-PD) cohort^22^, which was performed independently and with a different technology. Finally, we provide insights on how the key non-coding RNAs regulate gene expression by utilizing the long RNA sequencing data. Specific analyses on the long RNA-seq data are described in detail in a companion manuscript.

**Figure 1.**
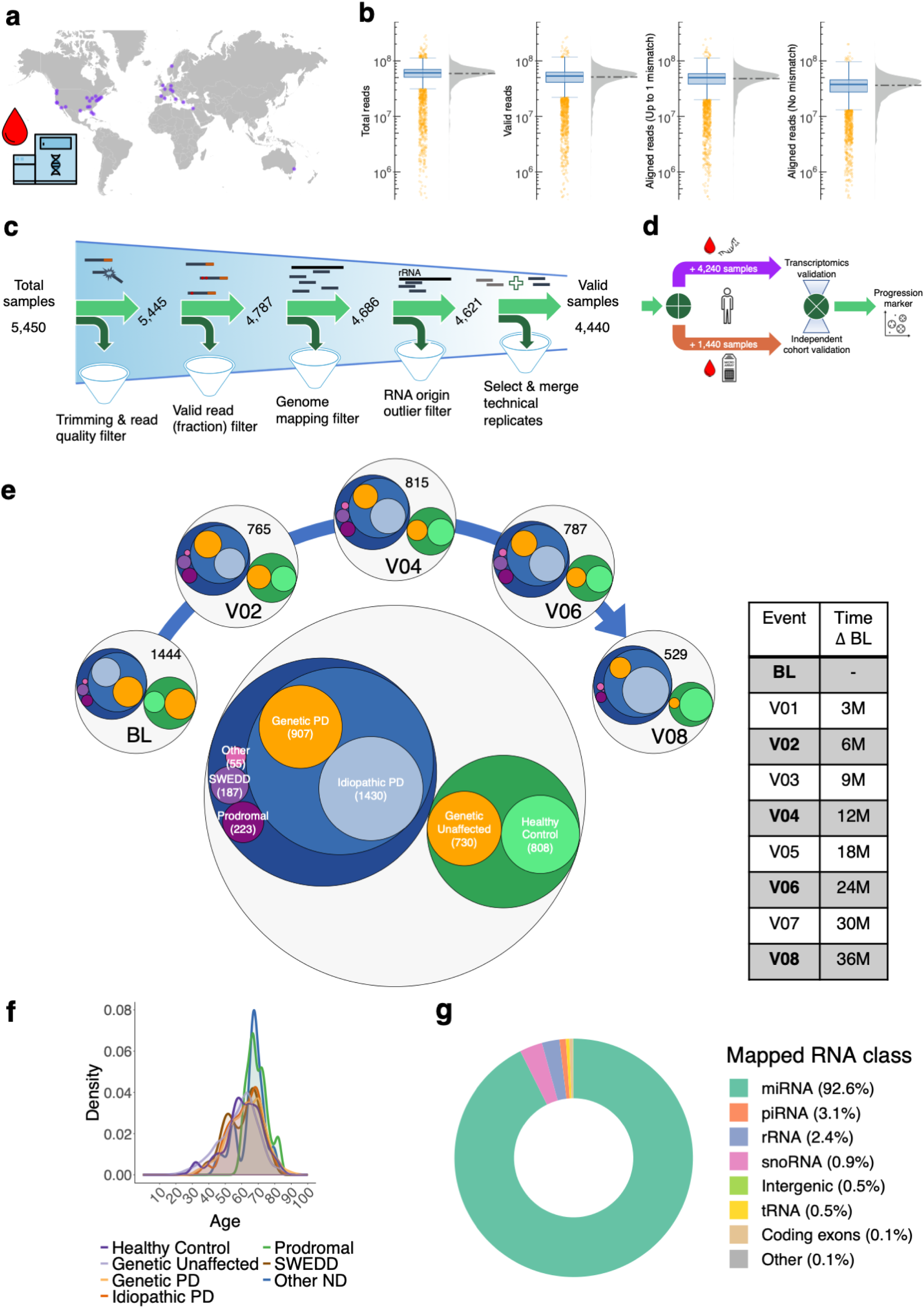
Study overview and sample characteristics. **a)** The PPMI project is a world-wide effort with 33 clinical sites where whole-blood samples of patients and healthy control donors were initially drawn. **b)** After RNA extraction, library preparation and sequencing by a single-vendor (Hudson-alpha), the sncRNA-samples exhibit a high number of reads and a good quality in general. Summary statistics as mean and standard deviation from left to right are: 59, 085, 191(± 31, 586, 910), 51, 112, 326(± 29, 542, 682), 47, 826, 485(± 27, 836, 420), and 36267825(± 21, 857, 435). **c)** Several stringent filter criteria were applied to each sequencing sample in order to remove potential outliers, after which approximately 80% of all samples remained for down-stream analysis. **d)** Further validation was performed using 1) paired mRNA sequencing data and 2) whole-blood microarray miRNA samples from the independent NCER-PD cohort. **e)** Distribution of valid samples according to **c)** across the investigated biogroups and timepoints (clinical visits) considered in this study. **f)** Age density distribution of cohort across the biogroups. **g)** Separation of valid sncRNA-seq reads into their genome origins identifies human miRNA precursors as predominant source. Shown are mean values with the standard deviation for each class being as follows: miRNA (±2.5), piRNA (±1.1), rRNA (±1.5), snoRNA (±0.5), intergenic (±0.2), tRNA (±0.1), coding exons (±0.02), and other (±0.004).

## Results

### High quality and ultra deep sncRNA-sequencing covers all major small RNA species

A known factor that affects sncRNA sequencing results severely is the sample integrity^26, 27^, calling for stringent quality control from the input to the sequencing results. The median RNA integrity number (RIN) of 8.2 points at an excellent RNA quality (Extended Data Figure 2a). Sequencing yielded a total of 322.6 billion reads at an average of 59 million per sample (Figure 1b, Extended Data Figure 1a). Of these 279 million (86%) passed a stringent quality control (Extended Data Figure 1b). Next, 93% of reads mapped to the human genome (Extended Data Figure 1c). Besides RNA input- and read quality control, the initial 5, 450 sncRNA-seq samples underwent a sample-wise quality control (Figure 1c). After sequential filtering, 4, 440 samples from 1, 511 individuals remained as high quality data set. Besides this high-throughput data set, we additionally generated two other large-scale data sets in order to validate findings and to understand downstream regulatory effects (Figure 1d). First, we sequenced the 4, 240 transcriptomics (RNA-seq) samples paired to the small RNA libraries from PPMI in order to assess potential targets of miRNAs (see details in the companion manuscript). Second, we validated key findings in an independent cohort of 1, 440 whole-blood miRNA-microarray samples. The curated high-quality sncRNA data set along with the validation in a second cohort and the long RNA sequencing of the same individuals facilitates the identification of diagnostic and prognostic PD biomarkers.

The PPMI samples comprise healthy controls, idiopathic (sporadic) cases of PD, a set of mutation carriers that is partitioned into genetic PD and unaffected individuals, prodromal subjects with isolated rapid eye movement behaviour disorder (RBD) and/or hyposmia, subjects showing scans without evidence of dopaminergic deficits (SWEDD), and other neurodegenerative diseases with similar symptoms (Figure 1e, Supplementary Table 1). In addition to samples collected at study baseline (BL), we included up to four additional samples (visits V02, V04, V06, and V08). Relative sub-cohort sizes show only slight differences at the different study time points. Further, cohort subjects are well-matched with respect to age (Figure 1f). The overall mean age being 61 (± 10.8) years in all samples, 62 (± 10.2) years mean age in total PD, 59 (± 11.6) years in controls and similar distributions in the remaining sub-cohorts.

In a first analysis we examined the distribution of reads to different RNA classes. With our original focus on miRNAs and optimised protocols for miRNA sequencing we reached the expected enrichment of this class. Over 90% of all reads belong to known miRNAs from miRBase v22. The very high read depth however also facilitates valid conclusions from other RNA species (piwi-interacting RNAs (piRNAs) (3.1%), ribosomal RNAs (2.4%), small nucleolar RNAs (snoRNAs) (0.9%), unspecific intergenic sequences (0.5%), transfer RNAs (tRNAs) (0.5%), coding exons (0.1%), and others (0.1%)). We did not observe significant shifts in the composition of the RNA classes between the disease and control groups (Extended Data Figure 1d-j). Moreover, samples and technical replicates showed very good correlation in terms of both sncRNA and miRNA counts (Extended Data Figure 2b,c). Also, principal components analysis (PCA) and batch effect assessment did not reveal an apparent bias in the data, although total sncRNAs appear to be slightly more affected by technical variables than just the subset of miRNAs (Extended Data Figure 2d-j). Having an initial picture of the data quality and the overall distribution of reads into the different RNA classes, we investigated molecular differences between the disease groups and controls.

### Molecular effects vary between different types of PD

The UMAP analysis of normalized miRNA expression produces a homogeneous but undifferentiated molecular picture on the samples, again not revealing obvious batch effects (Figure 2a). To find global differences between PD and controls we computed the density distribution of samples from each sub-cohort (Figure 2b). While samples from healthy controls and idiopathic cases of PD seem to form a larger cluster, patients and unaffected controls with genetic predisposition constitute another cluster. Interestingly, the prodromal cohort approximates the genetic cluster, while the samples from the SWEDD cohort are scattered. While biological variables outweigh the technical ones in a PVCA, a large fraction of data variance is not explained by available annotations (Extended Data Figure 2i,j).

**Figure 2.**
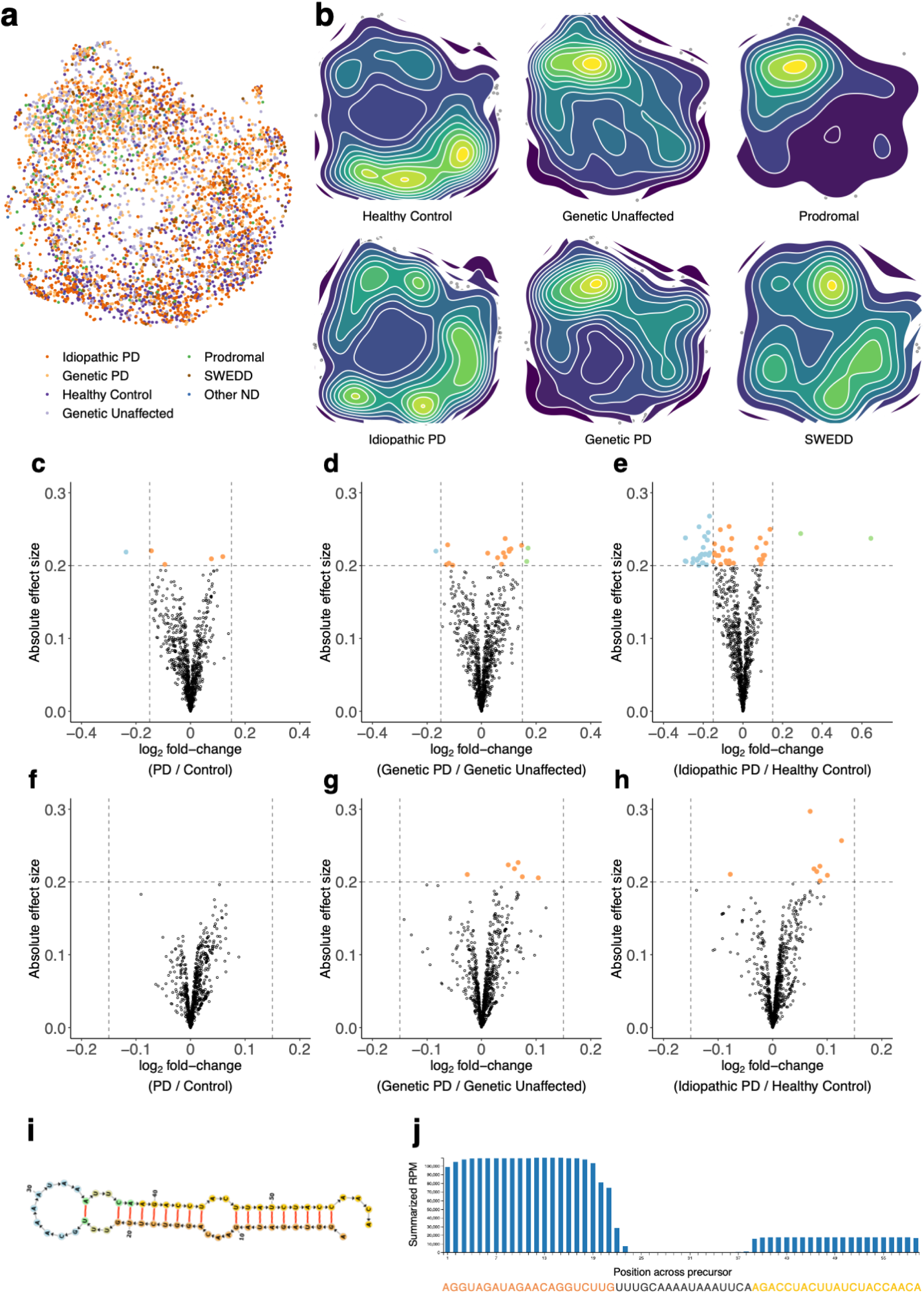
Association between miRNAs and Parkinson’s disease. **a)** Disease status colored UMAP embedding of all valid samples using the normalized human miRNA counts as features. **b)** Sample density plots according to UMAP embedding from **a)** separated by disease status. **c-h)** Volcano plots with (absolute) effect size (cohen s d) of expressed miRNAs for the main group comparisons considered in the PPMI study. MiRNAs that exhibit both a considerable effect size and fold-change are colored in blue or green when being up- or down-regulated, respectively. Orange points depict miRNAs with a considerable effect size but a small fold-change. **i)** Predicted precursor secondary structure of a novel miRNA for which both the −3p and −5p form exhibit a considerable effect size between total PD and controls. **j)** Mapped read profile for the novel precursor shown in **i)**. The −5p form is highlighted by dark orange sequence characters, while the −3p form is colored in yellow. Blue bars represent the sequencing coverage by the number of reads mapping to the precursor sequence.

We next investigated differential expression by phenotype and compared between PD (idiopathic + genetic) and controls (healthy + unaffected), the genetic cohort only, and patients without a genetic predisposition (Figure 2c-e). For the first comparison, we identified five miRNAs with a considerable effect size (miR-487b-3p: −0.22; miR-493-5p: −0.2; miR-6836-3p: 0.2; miR-6777-3p: 0.21 and miR-15b-5p: −0.21) in PD. The volcano plot indicates a trend towards a global down-regulation of miRNA abundance in PD. Remarkably, the comparison for the genetic cohort does not follow this global shift. In total, three miRNAs attract attention by their effect size, two of which are up-regulated in affected genetic carriers (miR-103a-2-5p: 0.22, *p* = 9.4 *×* 10^−4^; miR-339-5p: 0.22, *p* = 7.5 *×* 10^−4^) and one down-regulated (miR-15b-5p: −0.19, *p* = 0.0011). The third comparison suggests the non-genetic cohort as driver for the global down-regulation. While 23 miRNAs are down-regulated only two are up-regulated (miR-125b-5p: 0.24, *p* = 6.4 *×* 10^−6^, miR-100-5p: 0.23, *p* = 5.4 *×* 10^−5^). Overall, the effects on miRNAs seem to be more diverse and stronger in the sporadic cases of the disease than for the genetics (Supplementary Table 2).

The remarkable depth of the sequencing data facilitates calling of novel miRNA candidates. From an initial set of 30, 924 candidates reported by miRMaster^28^, 834 were manifested in a consistently detected readout across the sub-cohorts. For these high-profile candidates we repeated the differential expression analysis (Figure 2f-h). Novel miRNAs differ from known ones in two aspects. First, the differential expression shift seems to be flipped but consistent for the novel miRNAs in total PD and idiopathic PD. Second, hypothesis testing for the difference in expression resulted in 168, 51, and 110 significant miRNA candidates (adj. Wilcoxon p-values at *α* = 0.05) (Supplementary Table 3). Among the down-regulated candidates in idiopathic PD, we found nov-pred-mir-3679, showing a sufficient minimum free energy of −22 *kcal/mol* and a favourable precursor secondary structure (Figure 2i). The stem loop of the novel mir originates from chromosome 15 and exhibits an excellent read profile at both mature forms (Figure 2j), both of which were down-regulated in idiopathic PD. We also performed differential expression analysis by the other sub-cohorts and the patient gender and discovered both known and novel miRs to be de-regulated (Extended Data Figure 3a-j, Supplementary Tables 4,5). Analogous results for the remaining non-coding RNA classes are available in the supplement (Supplementary Tables 6,7). These results already suggest a substantial diagnostic potential of sncRNAs for PD. The next key question is whether miRNAs change also over time and with disease progression.

### Effects within PD depend on time, disease progression, and blood cell-types

We tested whether miRNAs show increasing effects over time and disease progression by analyzing the up to four follow-up visits for the PD patients. To this end, we performed differential expression analysis for all pairwise comparisons of clinical visits, i.e. BL vs V02, BL vs. V04, and so fourth. Remarkably, we discovered not only many miRNAs to be down-regulated within PD over time but observed an increasing trend correlating to the time that passed between visits (Figure 3a-j).

**Figure 3.**
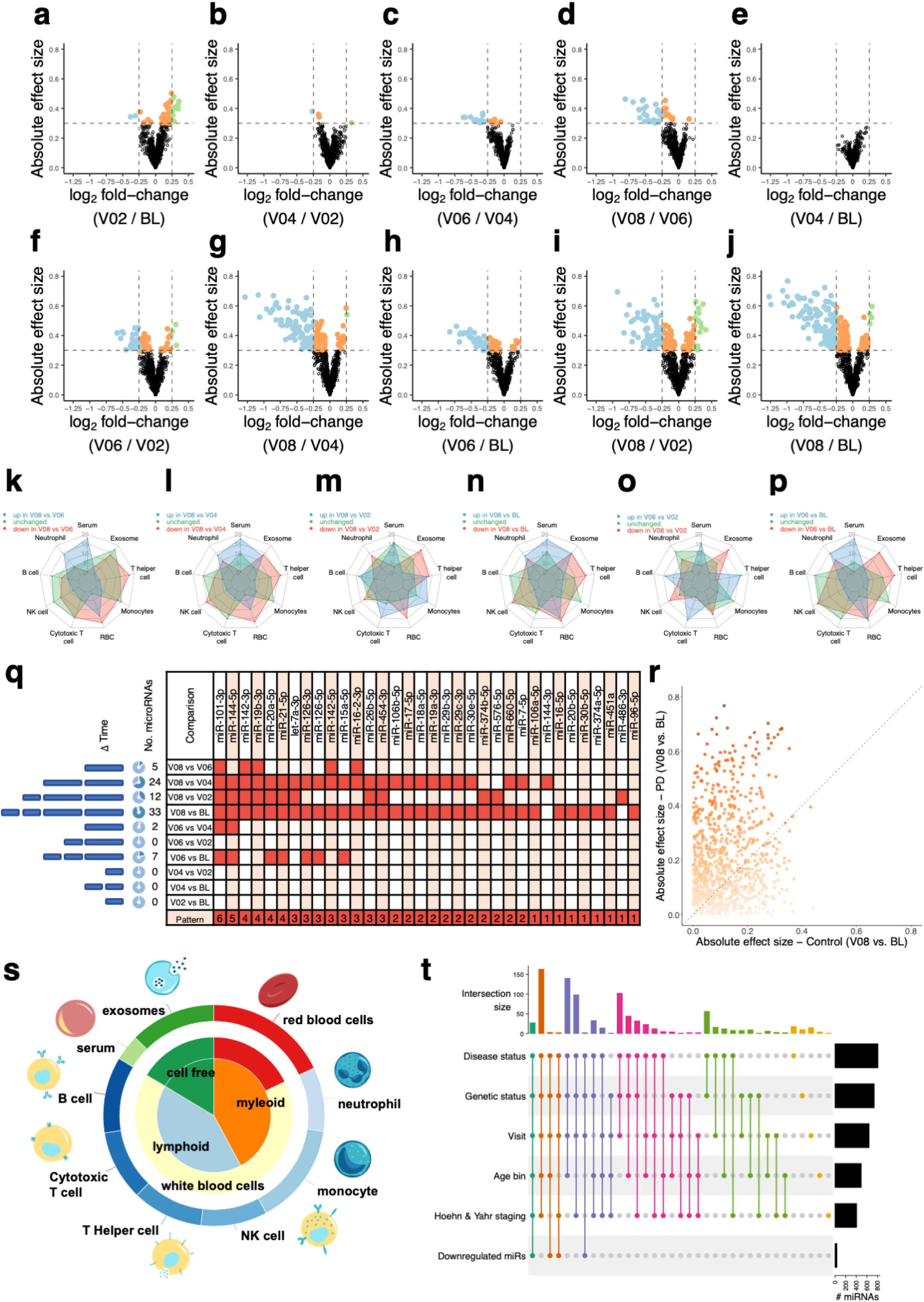
Progressive de-regulation of miRNAs within PD and predicted cell-type origins. **a-j)** Volcano plots with effect size of differential miRNA expression from the total PD samples for each pairwise comparison of the five clinical visits considered in this study. **k-p)** Six out of the ten pairwise comparisons of total PD samples yield de-regulated miRNAs, which show an uneven and varying distribution across the predicted cell-types of origin. **q)** Intersection of down-regulated miRNAs from a-j) that show a moderate effect size along the pairwise comparisons. Table cells colored in red denote a down-regulation in the respective combination of miRNA and timepoint comparison. The observed correlation with time passed between visits is indicated on the left. Small blue bars represent a timeframe of 6 months, while the large blue bars represent 12 months. **r)** Scatterplot showing absolute effect size for each miRNA comparing visits with the largest difference in time, i.e. latest visit (V08) versus baseline (BL), between total PD and total control. **s)** Relative composition of blood cell-type origins of down-regulated miRNAs displayed in **q). t)** Upset diagram showing all non-zero intersections of sets of significant miRNAs resulting from one-way ANOVAs for the sample annotation variables shown on the y-axis. The number of significant miRNAs per variable is shown on the right. The intersection size for each combination is displayed on top and colored according to the degree, i.e. the number of annotation variables intersected.

Since the cellular origin of these patterns has a potential impact, we deconvolved the blood-cell components and cell-type origins for both the de-regulated and unchanged miRNAs using the comparisons with the highest time delta (Figure 3k-p). We observe that down-regulated miRNAs mainly originate from exosomes, T helper cells, and red blood cells (RBCs) while up-regulated miRNAs are often associated to neutrophils and serum. These findings are consistent with an observed increase in neutrophils and decrease in lymphocytes identified in the PPMI blood cell count data, and further confirmed by transcriptomic changes in neutrophil-associated genes and pathways observed in the long RNA seq (details in the companion manuscript). Lastly, larger fractions of unchanged miRNAs were associated to B cells, NK cells, and cytotoxic T cells. For a systematic analysis of the observed depletion trends, we intersected the sets of miRNAs from all comparisons that exceed a fold-change threshold, resulting in 34 miRNAs, all of which are down-regulated in one to six of the 10 total comparisons (Figure 3q). Thereby, miR-101-3p and miR-144-5p showed the most consistently detected down-regulation (Extended Data Figure 3k). We then repeated the pairwise analysis scheme for the controls and not only observed significantly smaller effects than for PD, in particular for the comparison with the largest time difference (Figure 3r), but the magnitude of these differences again positively correlated with the number of miRNAs being consistent in each of the comparisons. Also, performing the blood cell-type deconvolution for the 34 miRNAs, we observed RBCs, exosomes, and monocytes to represent a large fraction, whereas only a minor fraction is associated to serum (Figure 3s). Furthermore, the analysis suggests the pool of leukocytes as a primary source of time-dependent depleted miRNAs in blood of PD patients.

For a rather global analysis of effects, we carried out an analysis of variance (ANOVA) of the miRNA expression for individuals across the disease groups, genetic status, clinical visits, age binnings, and Hoehn and Yahr staging. Comparing the sets of several hundreds of significant miRNAs obtained (Figure 3t), the largest pool of miRNAs is shared between all comparisons. In addition, the much smaller set of down-regulated miRNAs (Figure 3q) reveals a substantial concordance with the ANOVA results (Supplementary Table 8). These robust findings do not only suggest that sncRNAs have diagnostic and prognostic potential for PD but also indicate general age-dependent variability of miRNA biomarkers.

### PD shows two molecular ages of onset and patterns are highly age dependent

The PPMI cohort covers most of the average human lifespan and we leveraged miRNA expression to test whether molecular ages of disease onset exist. Indeed, the number of miRNAs de-regulated in PD is highly age-dependent and shows two ages of onset in the 30s and again beginning with the late 60s (Figure 4a). The number of differentially expressed miRNAs constantly declines in both idiopathic and genetic PD. We found a similar pattern for the SWEDD patients. Prodromal patients in general seem to have a late molecular onset and the number of de-regulated miRNAs increases with age. Surprisingly, the molecular miRNA landscape shows different traits when the directionality of regulation is considered (Figure 4b,c). The trends suggest a more severe down-regulation of miRNAs at the different ages of onset. Also, idiopathic and genetic PD patients show very distinct courses at early age for these miRNAs, while up-regulated miRNAs seem to behave vice-versa.

**Figure 4.**
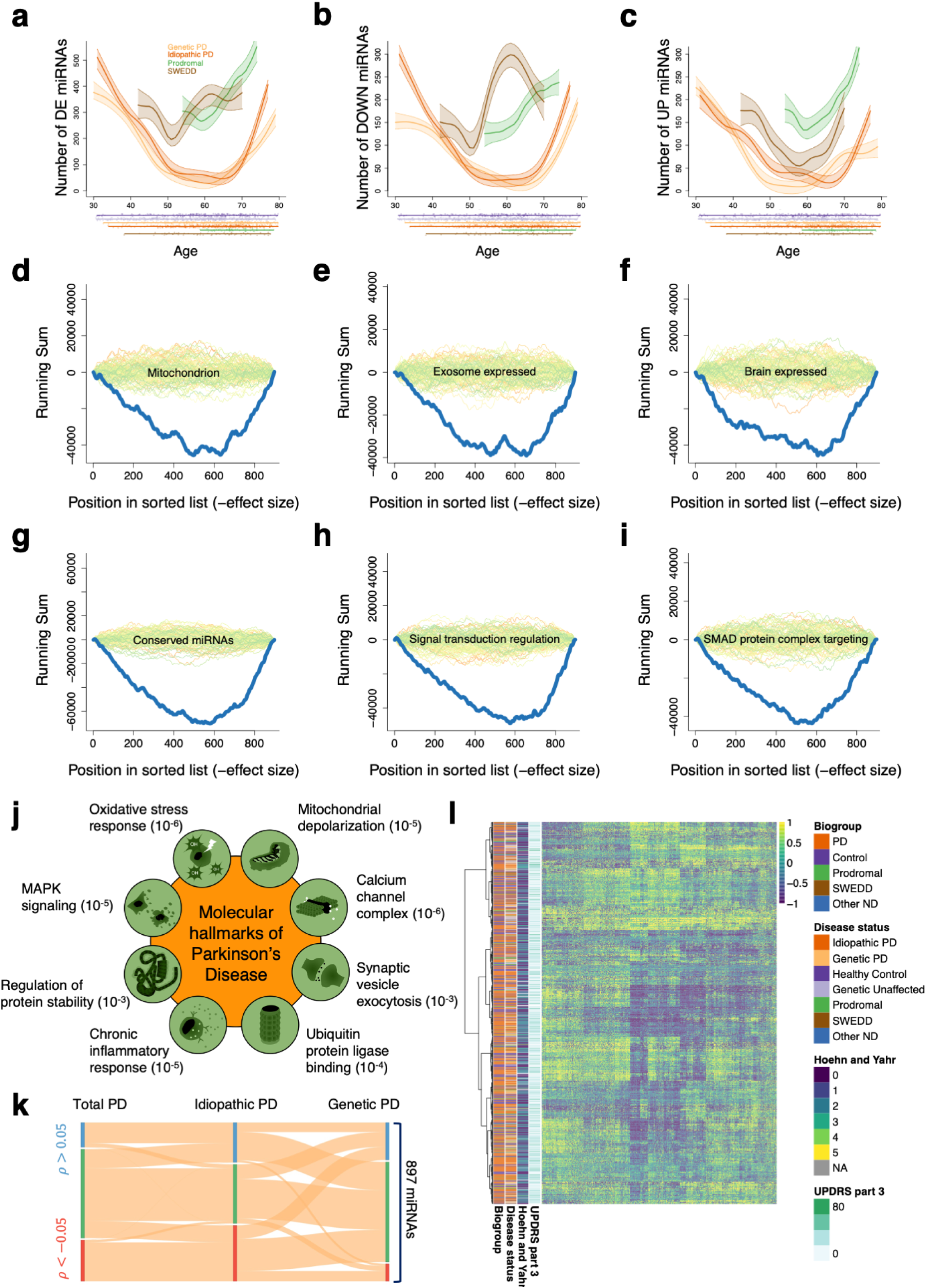
Age-related changes of miRNA expression and their association to the hallmarks of Parkinson’s highlight clusters for disease sub-types and progression. **a)** Spline smoothed number of differentially expressed miRNAs by a function of age of sample donor and by disease status. Dark lines depict the smoothed distribution while light colored bands centered around the curves correspond to the 95% confidence intervals. One-dimensional histograms at the bottom show the group-wise age distribution, including healthy controls (dark purple) and unaffected genetic carriers (light purple). **b)** Analogous to **a)** and limited to down-regulated miRNAs. **c)** Analogous to **a)** and limited to up-regulated miRNAs. **d-f)** Examples of significantly depleted biological categories and pathways observed from the list of miRNAs decreasingly sorted by effect size between total PD and control. Blue lines display the actual enrichment observed while green to orange lines show random enrichments (background) of simulated permutation experiments. **g-i)** Analogous to **d-f)** using miRNA effect sizes from the comparison of idiopathic PD vs healthy controls as input for enrichment analysis. **j)** Significantly de-regulated miRNA categories / pathways resemble the molecular hallmarks of Parkinson’s disease. Labels beside hallmark illustrations exemplify the categories as observed from the entire list of pathway enrichments/depletions along with a numeric scale for the adjusted p-values. **k)** Assignment of miRNAs according to their direction and degree of correlation with clinical visits (time), using the total PD samples (left), idiopathic PD only (middle), and genetic PD only (right). MiRNAs were classified as either being positively (blue), negatively (red), or not correlated (green) with time. Orange ribbons visualize the relative amount of miRNAs assigned from left to right and according to the correlation with time per disease group. **l)** Complete matrix of patient-wise spearman correlation between normalized expression of detected miRNAs and clinical visits. Patient (row) orders were obtained by a hierarchical clustering of the rows, annotated as dendrogram on the left. Major sample and patient annotations are also shown on the left of the heatmap. Likewise, miRNA (column) orders were obtained by hierarchical clustering (dendrogram omitted).

### Validation in an independent cohort

In the previous sections we described diagnostic and age dependent PD signatures. Although samples of many sites are included, biomarker discovery calls for a validation in an independent cohort and a different technology. We thus analyzed the PD cohort of the Luxembourg Parkinson’s Study, which comprises 1, 440 whole-blood derived microarray samples from 988 donors, each with up to four clinical visits (Supplementary Table 9). Of the participants 440 were diagnosed with idiopathic PD, 81 with Parkinsonism (atypical forms of PD), and 485 are age-matched healthy controls. Among the Parkinsonism sub-cohort, there are cases of Progressive Supranuclear Palsy (25), unspecified Parkinsonism (13), Cerebrovascular Disease with Parkinsonism features (13), Multiple System Atrophy (10), Lewy Body Dementia (10), Cortical-basal Syndrome (7), and Drug-Induced Parkinsonism (3). In total, 640 miRNAs passed QC (Supplementary Table 10) and 416 of those overlapped with miRNAs detected in the PPMI cohort. We carried out differential expression analysis between iPD and controls from NCER-PD and compared this to the effect sizes obtained for the three different comparisons in the PPMI cohort. Indeed, highest correlation was computed for the same comparison in the PPMI cohort (iPD vs. healthy control; Spearman correlation of 0.43). In contrast, the correlation between NCER-PD and PPMI for genetic PD vs. genetic unaffected decreased as expected to 0.21 (Extended Data Figure 4a-c). To find the biomarker set with highest concordance between NCER-PD and PPMI we performed a DynaVenn^29^ analysis, suggesting a highly significant overlap (adj. *p* = 10 *×* 10^−18^) of 222 miRNAs (Figure 4d-f).

We also repeated the analysis of differentially expressed miRNAs along the age axis for NCER-PD and observed a remarkable similarity to the PPMI cohort. Effect size of de-regulated miRNAs is highest in the third decade and seventh decade of life and reaches its minimum around the age of 65. We did not observe much difference between the number of down-regulated and up-regulated miRNAs, however the former show slightly larger effects at higher ages. Although very similar to the patterns for SWEDD, patients diagnosed with Parkinsonism exhibit three (≈ 50y, ≈ 62y, ≈ 80y) and two (≈ 62y, ≈ 80y) peaks for the number of down-regulated and up-regulated miRNAs, respectively. The clear diagnostic patterns that are largely consistent between the two cohorts open the question whether the biomarkers share similar molecular functions.

### De-regulated miRNAs resemble Hallmarks of Parkinson’s disease

Among the most significant hits in a miEAA^30^ pathway analysis, the down-regulated miRNAs in PD show a strong enrichment in the mitochondrion (Figure 4d, *p* = 9.57 *×* 10^−13^). Similarly, down-regulated miRNAs are enriched in exosome (Figure 4e, *p* = 4.34 *×* 10^−11^), microvesicle (*p* = 4.34 *×* 10^−11^) and are known to be freely circulating (*p* = 9.54 *×* 10^−9^). These findings also agree with the significant categories found by an over-enrichment analysis of the miRNAs validated in the NCER-PD cohort (Extended Data Figure 4g). Among the potential tissues of origin the top four are brain (Figure 4f, *p* = 1.10 *×* 10^−9^), spinal cord (*p* = 1.59 *×* 10^−8^), arachnoid mater (*p* = 1.19 *×* 10^−7^), and dura mater (*p* = 8.89 *×* 10^−7^). Our enrichment analysis suggested almost 200 disease hits with the most significant category of Alzheimer’s disease miRNAs. Among the top 10 we also observed neurodegenerative disease miRNAs in general (*p* = 1.17 *×* 10^−9^), arguing for a broader miRNA signature of related neurodegenerative disorders. For biological processes *regulation of T-helper cell differentiation* (*p* = 1.36 *×* 10^−6^), *regulation of neurotransmitter uptake* (*p* = 1.36 *×* 10^−6^), and *negative regulation of receptor signaling pathway via STAT* (*p* = 4.8 *×* 10^−6^) are most significant. Focusing on idiopathic PD vs. healthy controls and genetic PD vs. genetic unaffected resulted in two order of magnitudes more significant categories with substantially smaller p-values. For example, an observed depletion of vascular disease miRNAs (*p* = 4.02 *×* 10^−30^), conserved miRNAs (Figure 4g) and cellular signalling (Figure 4h-i) is exclusive to miRNAs de-regulated between idiopathic PD and healthy controls. Conversely, we found no significant pathways exclusively for patients from the genetic sub-cohort. Based on the enrichment analysis of total PD and controls we assessed the ability of blood-sampled miRNAs to resemble previously characterized molecular traits of PD. Indeed, we recovered a highly significant depletion of very specific molecular functions and pathways for each of the molecular hallmarks^31^ of the disease (Figure 4j).

The prominent pathway effects indicate distinct molecular regulatory patterns and open the question whether miRNAs also have a prognostic potential. In correlating miRNA expression with time and clinical events we confirmed the overall trend of decreasing miRNAs (Figure 4k) with strongest effect in iPD patients. A clustering of the matrix containing patient-wise spearman correlation values between miRNA expression and time identifies several clusters by disease group and Hoehn and Yahr staging (Figure 4l). The molecular patterns are in line with the observation that the pace of disease progression varies substantially between PD patients and usually worsens with higher ages, calling for a more detailed analysis of miRNAs in the context of individual disease progression.

### miRNAs are progression markers and constitute progression networks with targeted genes

According to clinical assessments of the Hoehn and Yahr staging, 68 patients switched to a lower staging level (non-progressing) with a difference in mean UPDRS part III score of −5.22 (V08 vs. BL), 601 slowly progressing patients remained in the same stage but showed a respective difference of +4.62, and 181 are fast progressing patients with a higher staging level at later visits and with difference in means of the score of +12.77 (Figure 5a). A correlation analysis of miRNA expression per PD patient from PPMI with at-least three visits suggested 71 miRNAs that are positively correlated in progressing patients while negatively correlated in non-progressing patients (Figure 5b). Vice versa, we observed 71 miRNAs to be negatively correlated with time in progressing cases of PD and positively in non-progressing individuals, providing further evidence for a prognostic potential. Testing another hypothesis we asked whether miRNAs and target genes are orchestrated in prognostic modules. To this end, we extended the correlation analysis to the whole transcriptome data of the same patients (details on this data set are available in the companion manuscript).

**Figure 5.**
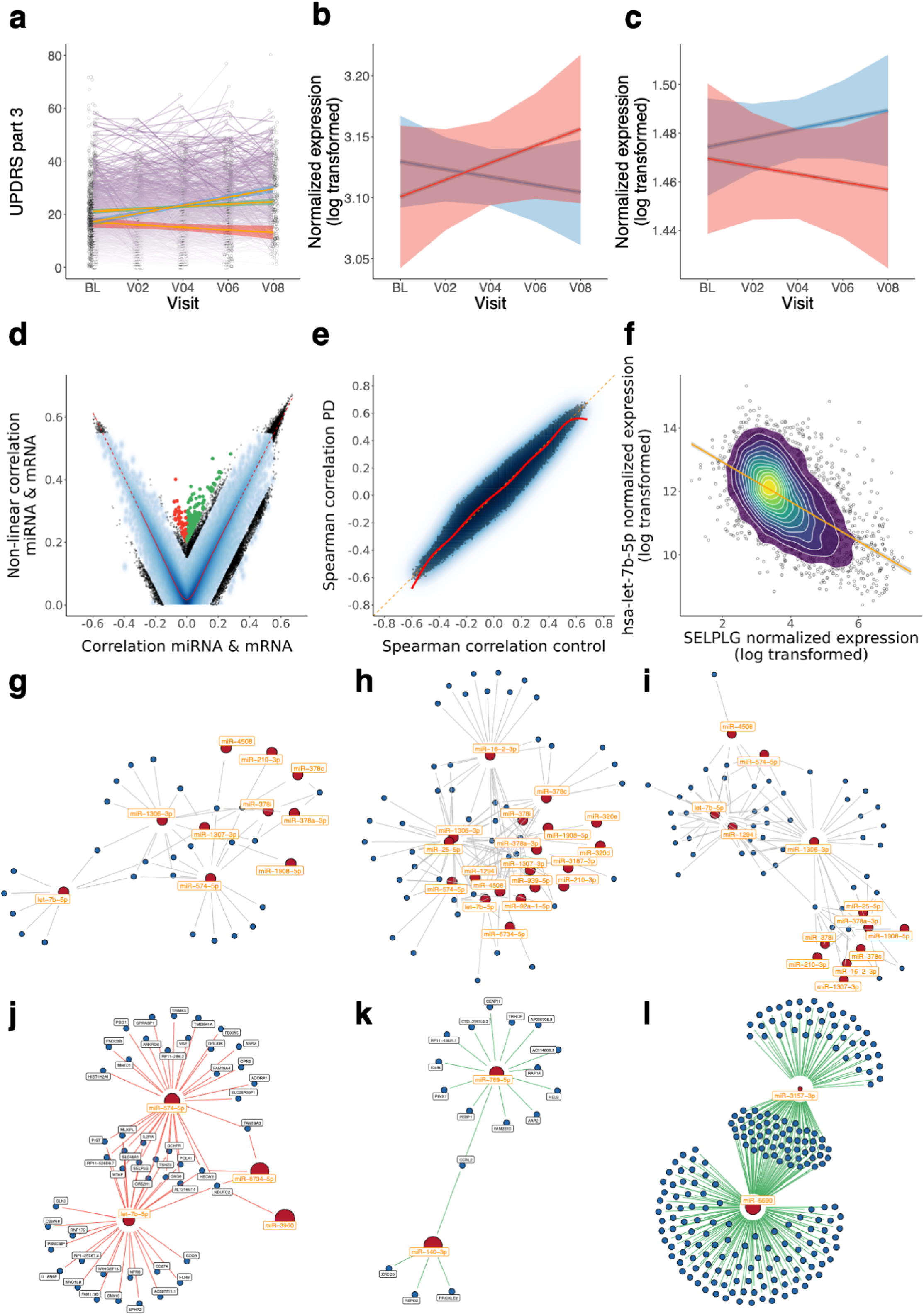
Progression marker analysis and down-stream validation using anti-correlated mRNAs. **a)** Distribution of disease progression in PD patients by clinical visit and UPDRS part 3 scoring. Patients were classified as either progressing (red trendline), non-progressing with the same Hoehn and Yahr staging in the last visit as for baseline (green trendline), or showing ameliorated symptoms with a lower staging as compared to baseline (blue trendline). Light bands around the three trendlines correspond to the 95% confidence intervals. **b)** Smoothed trendline of expression levels for miRNAs showing a positive correlation with clinical visits in progressing patients (red) vs. non-progressing patients (blue). Light bands around the trendlines mark the 95% confidence intervals. **c)** Analogous to **b)** but for miRNAs exhibiting a negative correlation with clinical visits. **d)** Linear vs. non-linear correlation of paired miRNA and mRNA samples across all biogroups. **e)** Spearman correlation of paired miRNA and mRNA normalized expression scattered between total PD samples and controls. **f)** Case example of negative correlation between normalized expression levels of hsa-let-7b-5p and SELPLG. **g-i)** Inferred miRNA-mRNA regulatory networks as bipartite graphs of the top-correlated modules in pooled controls, idiopathic PD, and genetic PD. Red nodes represent miRNAs and blue nodes mRNAs. Pairs of miRNAs and mRNAs are connected if the inverse correlation exceeds a pre-set threshold (cf. Materials and methods). **j)** Example miRNA-mRNA regulatory module using only samples from disease progressing patients. Orange colored miRNAs are negatively correlated with both disease progression and potential mRNA targets as extracted from the inferred networks. Radii of miRNA nodes are scaled relatively to the degree of progression. **k**,**l)** Similar to **j)** where miRNAs exhibit a positive correlation with disease progression, while preserving a negative correlation with potential targets.

Comparing the linear and non-linear correlations of all pairs of miRNA and mRNA, we observed pre-dominantly a linear dependency, although some exceptions exist (Figure 5d). The correlation of miRNAs and mRNAs in PD patients versus controls resulted in a high agreement but also small deviations (Figure 5e). Intriguingly, some of the miRNAs are almost perfectly anti-correlated to potential targets (Figure 5f). Since the interactions between miRNAs and mRNAs can be unspecific when viewed individually, we assumed a *M* : *N* relationship by constructing bi-partite regulatory graphs. Based on all anti-correlated pairs we constructed putative miRNA-mRNA core networks for pooled controls, idiopathic PD, and genetic PD that exceed the same edge threshold (Figure 5g-i). Surprisingly, we observed a difference in terms of number of nodes and edge density between the three groups of samples. Both disease-derived networks display an increased number of miRNAs and targets with some nodes being shared, e.g. let-7b-5p and some being exclusive to the disease, as for example miR-25-5p. Examining possible changes of the networks in relation to disease progression required a re-computing of graph components for the down-regulated miRNAs for which putative targets are up-regulated in progressing PD and vice versa. Thereby, we identified three core components. First, up-regulated targets in progressive PD patients are potentially regulated by four down-regulated miRNAs (Figure 5j). Two more subnetworks with two up-regulated miRNAs, hsa-miR-769-5p and hsa-miR-140-3p, as well as hsa-miR-3157-3p and hsa-miR-5690, showed down-regulated and partially shared target genes (Figure 5k,l).

## Discussion

As the world population ages and the mean life-expectancy increases, neurodegenerative disorders show a steadily rising incidence. To counteract these trends, large-scale disease progression studies are of immense value towards the development of curative treatments. The PPMI has accumulated a high-quality data set of small-RNAs sequenced from whole-blood of healthy controls and PD patients from multiple disease stages at an unprecedented scale. Our in-depth analysis of this data set highlights the complex changes of small-RNA, in particular miRNA, expression upon aging and disease onset.

While we generated data on basically all small RNA classes at high sequencing depth, we focus on the interpretation of microRNAs, which represents the majority of the reads. Given the cohort design and statistical power facilitated by PPMI, effects are considerable and highly significant. Surprisingly, miRNA expression signatures distinguish between iPD and genetic PD, for which effects between disease and controls were largest for iPD. The observed down-regulation of miR-15b-5p is in line with a proposed signature of 5 serum miRNAs in PD patients^32^ and a miRNA-mRNA interaction network analysis of PD progression^33^. We also observed a systematic decline in miRNA expression over time, revealing a signature of miRNAs associated with leukocytes, in particular monocytes and B cells, as well as exosomes and RBCs. Among these are several members of the miR-19 and miR-29 families that are consistent with earlier findings^33–37^. Also, PD induced changes for both monocyte and exosome expression signatures has been reported in several other studies^38–41^. Our results also highlighted a strong age related component of miRNA expression with two distinct waves of molecular onset. Ravanidis *et al* recently showed differentially expressed, brain-enriched miRNAs to have discriminatory power between iPD and genetic PD^42^. Comparably, we report a highly significant down-regulation of mitochondria, exosome, and brain expressed miRNAs over time in PD. In particular, mitochondria dysfunction seems to play a major role in the disease, although it is still unclear whether it’s part of the cause or a consequence of the disease^43^. Shamir *et al*^44^ described affected gene interactions networks in iPD to be associated with oxidation, ubiquitination and other hallmarks of PD, an effect that is mirrored by the dys-regulated miRNAs revealed in our analysis of the PPMI cohort.

Of interest are also the novel miRNA candidates. In this context, novel means that the miRNAs are either not annotated for *h. sapiens* in the latest miRBase release 22.1, or haven’t been described before. The novel miRNA candidate we report to be of relevance for PD is one example where miRNAs are annotated for model organisms but human. A miRBlast analysis using miRCarta^45^ revealed that this novel human miRNA is known in various other species, including *r. norvegicus* or *m. musculus* (rno-mir-1839 and mmu-mir-1839). In fact, this increases the likelihood that respective candidates are no artefacts but indeed genuine human miRNAs.

Large-scale biomarker discovery studies call for a validation in an independent cohort using an independent technology. In this regard, we observed a good concordance between NGS-samples from PPMI and microarray samples from the Luxemburg NCER-PD cohort. Still, challenges with these studies remain. The data set exhibits varying number of samples for the different PD types and staging schemes, further complicating the analysis. Likewise, the age range of the cohorts is not covered uniformly between 30 and 70 years, potentially influencing the age dependent results. Moreover, the number and the classification-scheme of distinct PD sub-types is still subject to on-going clinical research. We tried to address this uncertainty by comparing the sub-cohorts in several ways, once with pooled controls and the genetic and non-genetic cohorts independently. Further, the manifestation of RBD in prodromal has not yet been broadly characterised, however the absence of motor symptoms suggests an early disease-stage for these patients, who then show a late PD-onset, an effect that we confirm from a molecular viewpoint. In contrast, SWEDD patients stand out as they are not part of the other clinical continua and show similar but also distinct molecular patterns. Further investigation specifically in the direction of SWEDD is thus deemed necessary. There exist also confounding variables, for example, PD subjects were required to be drug-naïve at study enrollment but could possibly be treated with PD medications at any of the later clinical visits. Our consecutive comparison of all clinical visits nevertheless suggests a systematic and drug-independent effect induced upon the miRNAs that has been reported earlier^46^. Further, vague symptoms can lead to diagnostic uncertainty, a fact that is reflected by a small subset (*<* 10) of PD patients from PPMI who received a differential diagnosis at later clinical visits. However, given the design stringency and scale of the presented study we expect a high tolerance level towards such exceptions.

Another important factor are potential batch effects, for which we found no apparent bias in the data, however they are difficult to quantify in general and thus can remain hidden. The greatest challenge is to determine the specificity of affected miRNAs towards PD. As one example, miR-144 that we discovered as a prominent marker in the present study has already been described to be a general disease marker^47^. Also, the quite heterogeneous mixture of blood cells can skew the molecular patterns. Here, accurate single cell studies will certainly contribute to an improved resolution in the near future.

In summary, the here presented data set, which is one of the most complex if not the most comprehensive RNA-seq study on PD, is an excellent resource to conduct analyses of blood-borne biomarkers with great statistical power.

## Online Methods

### Cohort design and blood sample collection

Data used in the preparation of this article were obtained from the Parkinson’s Progression Markers Initiative (PPMI) database (https://www.ppmi-info.org/data). For up-to-date information on the study, visit https://www.ppmi-info.org/. 4,690 longitudinal patient blood samples from 1,614 individuals were collected as part of the Parkinson’s Progression Marker Initiative (PPMI). Subject blood samples were collected by venous draw during periodic clinical visits alongside clinical and imaging data at each longitudinal timepoint. Blood samples were captured in 2×2.5mL PAXgene Vacutainer tubes, mixed by serial inversion, and incubated at room temperature (18-25C) for 24 hours, and stored at −80C prior to purification. 1ug of RNA purified according to the Qiagen PAXgene blood miRNA kit protocol (cat:763134) was used for this analysis. A comprehensive study overview, complete sample collection schedules, and clinical protocols are available at https://www.ppmi-info.org/.

### smallRNA Library Preparation and sequencing for PPMI project

The concentration and integrity of the RNA were estimated using Quant-It RiboGreen RNA Assay Kit (Thermo Fisher Scientific) and High Sensitivity RNA kit on the 5300 Fragment analyzer (Agilent Technologies), respectively. Approximately 200 ng of Total RNA from each sample was taken into small RNA library preparation protocol using Automated NEXTflex® Small RNA-Seq Kit v3 (Bioo Scientific, PerkinElmer) for Illumina® Libraries on PerkinElmer® Scilone® G3 NGS workstation according to manufacturer’s protocol. Briefly, 3’ 4N adenylated adapters mix were ligated to total input RNA followed removal of excess Adapters using adapter inactivation buffers. Post Ligation purification was done 2 times using the NEXTflex Cleanup beads, the purified material was eluted in 10ul of Nuclease Free Water. 5’ 4N adapters were then ligated to the RNA samples. Reverse transcription (RT) was done using M-MuLV reverse transcriptase for 30 minutes at 42°C and then for 10 minutes at 90°C. After the RT step the samples were cooled at 4°C in the thermal cycler. Post RT samples were spun down at 2000 rpm for 2 minutes and then stored in −20°C freezer overnight. The following day, Post-RT material was purified using NEXTflex Cleanup beads, with elution in 22.5ul Nuclease Free Water into NEXTflex barcoded PCR primer Mix. PCR setup was done using NEXTflex Small RNA PCR Master Mix and the amplification was performed at 95°C for 2 minutes followed by 20 cycles of 95°C for 20 seconds, 60 for 30 seconds and 72 for 15 seconds, final elongation was done at 72 for 2 minutes. Post PCR dual size gel free size selection was done on the Scilone G3 using NEXTflex Cleanup Beads with final elution made in 15ul NEXtflex Resuspension buffer. From the Post-PCR purified final libraries, 2ul of each library was taken for quality check, a 2 x dilution plate was made and the final library concentration and profile were assessed using Quant-iT Picogreen dsDNA Assay Kit (Thermo Fisher Scientific) and High Sensitivity (HS) DNA Assay on the Caliper LabChip Gx (PerkinElmer Inc.), respectively. qPCR was performed on final libraries using KAPA Biosystems Library Quantification kit (Kapa Biosystems, Inc.) to determine the exact nano molar concentration of those. Each library was diluted to a final concentration of 1.5nM and pooled in equimolar ratios, Single End (SE) sequencing (50bp) was performed to generate approximately 50 million reads per sample on an Illumina NovaSeq 6000 sequencer (Illumina, Inc).

### RNA extraction and microarray experiments from NCER-PD

For quantification of RNA eluates, Nanodrop ND-1000 Instrument was used (Thermo Fisher Scientific, Darmstadt, Germany). Quality control was performed using Agilent 2100 Bioanalyzer according to the manufacturer’s instructions (Agilent Technologies, Santa Clara, CA, USA). The expression profiles of all miRBase release v21 human miRNAs were determined using Agilent Sureprint G3 Human miRNA (8 × 60 K) microarray slides. Each array targets 2,549 microRNAs with 20 replicates per probe. 300 ng total RNA was dephosphorylated, labelled and hybridized using the Agilent’s miRNA Complete Labeling and Hybridization Kit according to manufacturer’s protocol. After hybridization for 20 h at 55 °C, the slides were washed twice and scanned using Agilent’s High Resolution Microarray Dx Scanner. Scan images were transformed to raw text data using Feature Extraction Software v12.0.3.1 (Agilent Technologies, Santa Clara, CA, USA).

### Quality control and sample preprocessing

For the PPMI sncRNA-seq evaluation, BCL’s were converted to FASTQ using bcltofastq v1.8.4, and FASTQ’s were merged and processed with miRMaster v1.0^28^. Adapters with up to an edit distance of 1 and a minimum overlap of 10 nt were trimmed with miRMaster and the 4 random nucleotides at the 5’ end of the sequence as well as in front of the adapter were removed. No unknown bases were allowed and reads were quality trimmed at the 3’ end when the mean quality of a window of 4 bases dropped below 20. Reads were then aligned to the GRCh38 genome with Bowtie (1.1.2). To quantify miRNAs, reads were mapped to miRBase v22 precursors with Bowtie and processed with miRMaster to allow up to 1 mismatch and 2 nt overlap at the 5’ end and 5 nt overlap at the 3’ end of the miRNA annotation. Other sncRNAs were quantified by mapping reads to the Ensembl ncRNA 85 sequences, the tRNAs from GtRNAdb 2.0 and the piRNAs of piRBase 1.0. The overall processing to yield valid reads and valid samples, comprises read trimming and quality filtering, valid read fraction filtering, genome mapping distribution filtering, RNA origin outlier filtering, and merging of technical replicates. For the primary analysis, raw expression matrices of sncRNAs, miRNAs, and novel miRs were filtered to discard features with fewer than 5 raw reads in 50% of the samples in each sub-cohort (Genetic PD, Genetic Unaffected, Healthy Control, Idiopathic PD, Other ND, Prodromal, SWEDD). Remaining counts were rpm normalized and log_2_-transformed. For the comparative analysis of the PPMI and NCER-PD cohorts, counts were quantile-normalized and log_2_-transformed.

For the PPMI whole-transcriptome data, BCL’s were converted to FASTQ using bcltofastq v1.8.4, and FASTQ’s were merged and quantified with Salmon (v0.7.2) based on GRCh37(hs37d5) and GENCODE v19. More details on the preprocessing can be found in the transcriptomics companion manuscript. For the combined analysis of miRNA and mRNA, miRNA counts were normalized to reads per million mapped to miRNA (rpmmm) normalized and mRNAs normalized to transcripts per million (tpm), and both log_2_-transformed. During down-stream analysis only paired samples for each individual having a unique match of sequencing IDs were considered.

For the NCER-PD microarray evaluation, samples were processed as previously described^30^. Based on the computed sample detection matrix, features were filtered to exclude miRNAs with a detection rate less than 50% in each sub-cohort (Idiopathic PD, Parkinsonism, Control). Expression signals were quantile-normalized and log_2_-transformed. The entire sample preprocessing procedure was implemented in R v3.5.1 with data.table v1.12.0, bit64 v0.9.7, preprocesscore v1.46.0 libraries, and Python v3.6.7 with Numpy v1.16.4.

### Bioinformatics and data analysis

Down-stream analysis was performed using R with multiple software packages listed in the following. For differential expression analysis, log_2_ fold-changes based on normalized expression *e* were calculated as log_2_(*e*_*case*_*/e*_*control*_) and effect size was calculated with the function cohen.d from the effsize v0.7.6 package with arguments (*e*_*case*_, *e*_*control*_). Statistical tests (Student’s t-test, Wilcoxon rank-sum test, Shapiro–Wilk test, One-way anova / F-test, Kruskal–Wallis test) were performed two-tailed using the R stats implementations. Wherever applicable 95% confidence intervals were used. All plots shown were generated using either base R functionality and / or functions from the ggplot2 v3.2.1, hexbin v1.28.1, pheatmap v1.0.12, rcolorbrewer v1.1.2, viridis v0.5.1, cowplot v1.0.0, ggrepel v0.8.1, ggsci v2.9, gplots v3.0.1.2, ggpubr v0.2.4, highcharter v0.7.0, fmsb v0.7.0, gridgraphics v0.4.1, ggrastr v0.1.7, ggformula v0.9.2, ggextra v0.9, upsetr v1.4.0, and complexheatmap v2.2.0 packages. Dimension reduction by PCA was accomplished using the prcomp function from R stats and by UMAP^48^ using the corresponding umap v0.2.3.1 package. Network analyses were performed using igraph v1.2.4.2, ggraph v2.0.1, networkd3 v0.4, rgraphviz v2.30.0, and grbase v1.8.3.4. Common data manipulation tasks were implemented using data.table v1.12.8, openxlsx v4.1.4, scales v1.1.0, stringr v1.4.0, and rfast v1.9.5. Precursor secondary structures and read profiles were extracted from the web reports of miRMaster. For the blood component deconvolution of miRNAs, we utilised a miRNA *×* cell-type matrix from a previously published NGS data set (GSE100467)^49^. First, rows were transformed into percentages by dividing each row by its sum. Next, percentages across origins were obtained by dividing each column by its sum of the row percentages. Then, values were extracted and normalized for all miRNAs of interest. One-way ANOVA p-values were obtained using the aov function of the R stats package. P-values were corrected in any case using the false discovery rate (FDR)-controlling procedure by Benjamini-Hochberg, also implemented in R stats. Adjusted p-values smaller than 0.05 were considered significant unless stated otherwise. For the aging trajectories of de-regulated miRNAs, effect sizes were computed for binned ages at study consent values using a sliding window approach. For each integer *i* in the interval [30, 80], effect size (cohen’s d) was calculated for each miRNA using samples from the agebin [*i, i* + 10). Thereby, we required at least 10 samples from both case and control group and a minimum absolute value of 0.3 to consider the effect size for a miRNA at age *i*. The resulting binary matrix of dimension miRNA *×* age bins was summed up column-wise to yield a vector containing the number of miRNAs de-/up-/down-regulated at each age bin. Final trajectories were obtained by applying the function smooth.spline from R stats with the age bins as predictor variable, and the summarized counts as response variable while allowing eight degrees of freedom. The upper and lower 95% confidence bands were calculated using the jackknife residuals. Spearman and Pearson correlation coefficients were computed with the cor function of R stats. To perform Principal Variance Component Analysis (PVCA) to estimate any influence of batch variables on the expression and the degree of expression variance associated to the annotation variables, we used the function pvcaBatchAssess from the R package pvca v1.28.0. MiRNA set enrichment (Kolmogorov-Smirnow test) and over-representation analyses (Hypergeometric tests) were performed with miEAA 2.0^30^. Analysis of significant overlaps between miRNAs detected in either NGS or the microarray data was accomplished with DynaVenn^29^, once for the two lists of miRNAs sorted by increasing AUC and once sorted by decreasing AUC. Patients were classified as progressing if Hoehn and Yahr staging increased in the last available clinical visit according to baseline, and as non-progressing otherwise. Patient-wise spearman correlation of expression and time (clinical visits) was calculated and averaged over all patients for each miRNA. For the regulation network analysis we calculated Spearman’s *ρ* between paired miRNA and mRNA samples and investigated the subnetworks spawned by pairs exceeding a given threshold for anti-correlation, which was set to be at least −0.35. On a case-by-case analysis for the different group comparisons, thresholds on edge-values were further numerically decreased to yield smaller but more specific networks. On these subnetworks we searched for strongly connected components. For the progression networks, we computed the miRNA-mRNA correlations separately for progressing and non-progressing patients and kept miRNAs that had flipped signs of mean correlation with time in progressing vs. non-progressing patients. Next, we discarded miRNAs and mRNAs exhibiting a mean expression shift between first and last clinical visit of disease progressing patients smaller than 0.05. The final networks displayed were obtained analogously to as described above. The final manuscript figures were compiled using Microsoft’s PowerPoint v16.36.20041300.

## Acknowledgements

PPMI – a public-private partnership – is funded by the Michael J. Fox Foundation for Parkinson’s Research and funding partners, including Abbvie, Allergan, Amathus Therapeutics, Avid, Biogen, BioLegend, Bristol-Myers Squibb, Celgene, Denali, GE Healthcare, Genetech, GlaxoSmithKline, Handl Therapeutics, Insitro, Janssen Neuroscience, Lilly, Lundbeck, Merck, MSD, Pfizer, Piramal, Prevail, Roche, Sanofi Genzyme, Servier, Takeda, Teva, UCB, Verily, Voyager, and Golub Capital. We highly appreciate the encouragement and support of Karoly Nikolich in setting up and performing the study. The microarray experiments have been performed as fee-for-service by Hummingbird Diagnostics. We acknowledge the support of HbDx. We would like to give special thanks to all participating patients in the study. Additionally, we are very grateful for all received funding and private donations that enabled us to carry out the project. Furthermore, we acknowledge the joint effort of the NCER-PD consortium members generally contributing to the Luxembourg Parkinson’s Study: Aguayo, Gloria; Allen, Dominic; Ammerlann, Wim; Aurich, Maike; Baldini, Federico; Balling, Rudi; Banda, Peter; Beaumont, Katy; Becker, Regina; Berg, Daniela; Betsou, Fay; Binck, Sylvia; Bisdorff, Alexandre; Bobbili, Dheeraj; Brockmann, Kathrin; Calmes, Jessica; Castillo, Lorieza; Diederich, Nico; Dondelinger, Rene; Esteves, Daniela; Ferrand, Jean-Yves; Fleming, Ronan; Gantenbein, Manon; Gasser, Thomas; Gawron, Piotr; Geffers, Lars; Giarmana, Virginie; Glaab, Enrico; Gomes, Clarissa P.C.; Goncharenko, Nikolai; Graas, Jérôme; Graziano, Mariela; Groues, Valentin; Grünewald, Anne; Gu, Wei; Hammot, Gaël; Hanff, Anne-Marie; Hansen, Linda; Hansen, Maxime; Haraldsdöttir, Hulda; Heirendt, Laurent; Herbrink, Sylvia; Hertel, Johannes; Herzinger, Sascha; Heymann, Michael; Hiller, Karsten; Hipp, Geraldine; Hu, Michele; Huiart, Laetitia; Hundt, Alexander; Jacoby, Nadine; Jarosław, Jacek; Jaroz, Yohan; Kolber, Pierre; Krüger, Rejko; Kutzera, Joachim; Landoulsi, Zied; Larue, Catherine; Lentz, Roseline; Liepelt, Inga; Liszka, Robert; Longhino, Laura; Lorentz, Victoria; Mackay, Clare; Maetzler, Walter; Marcus, Katrin; Marques, Guilherme; Martens, Jan; Mathay, Conny; Matyjaszczyk, Piotr; May, Patrick; Meisch, Francoise; Menster, Myriam; Minelli, Maura; Mittelbronn, Michel; Mollenhauer, Brit; Mommaerts, Kathleen; Moreno, Carlos; Mühlschlegel, Friedrich; Nati, Romain; Nehrbass, Ulf; Nickels, Sarah; Nicolai, Beatrice; Nicolay, Jean-Paul; Noronha, Alberto; Oertel, Wolfgang; Ostaszewski, Marek; Pachchek, Sinthuja; Pauly, Claire; Pavelka, Lukas; Perquin, Magali; Reiter, Dorothea; Rosety, Isabel; Rump, Kirsten; Sandt, Estelle; Satagopam, Venkata; Schlesser, Marc; Schmitz, Sabine; Schmitz, Susanne; Schneider, Reinhard; Schwamborn, Jens; Schweicher, Alexandra; Stallinger, Christian; Simons, Janine; Stute, Lara; Thiele, Ines; Thinnes, Cyrille; Trefois, Christophe; Trezzi, Jean-Pierre; Vaillant, Michel; Vasco, Daniel; Vyas, Maharshi; Wade-Martins, Richard; Wilmes, Paul.

## Author contributions statement

FK led the statistical analysis of the data and contributed writing the manuscript; TF supported the statistical analysis of the data; IV and EA contributed to the primary analysis of data and matching to clinical variables; EH contributed to the general analysis of sequencing data; MK and CB assisted the general analysis of the microarray data; NLG supported the statistical analyses and visualisation of aggregated data. PG supported the statistical analysis with a focus on disease progression; KP contributed to the clinical interpretation of the data; BC contributed to study set up and interpretation of the data; RB, LG and RK contributed to the set up of NCER-PD and the interpretation of the microarray data; DG and BM contributed to the development of standard operating procedures, participant recruitment, and clinical interpretation of the data; EM contributed to the miRNA-gene interaction network analysis and the manuscript. TWC contributed in interpreting and discussing the data as well as to the manuscript. DWC and KKJ worked on the mRNA expression data and blood cell type analyses; AK contributed to study set up, supported the statistical analyses, and contributed to manuscript writing.

## Additional information

### Supplementary data

- **Supplementary Table 1:** Full sample and patient annotations for the PPMI cohort.
- **Supplementary Table 2:** Results of primary differential expression analysis comparisons of miRBase v22 miRNAs for the PPMI cohort.
- **Supplementary Table 3:** Results of primary differential expression analysis comparisons of novel miRNA candidates for the PPMI cohort.
- **Supplementary Table 4:** Results of secondary differential expression analysis comparisons of miRBase v22 miRNAs for the PPMI cohort.
- **Supplementary Table 5:** Results of secondary differential expression analysis comparisons of novel miRNA candidates for the PPMI cohort.
- **Supplementary Table 6:** Results of primary differential expression analysis comparisons of known sncRNAs for the PPMI cohort.
- **Supplementary Table 7:** Results of secondary differential expression analysis comparisons of known sncRNAs for the PPMI cohort.
- **Supplementary Table 8:** ANOVA results for primary sample annotation variables using miRBase miRNA counts from the PPMI cohort.
- **Supplementary Table 9:** Full sample and patient annotations for the NCER-PD cohort.
- **Supplementary Table 10:** Results of primary differential expression analysis comparisons of miRBase v21 miRNAs for the NCER-PD cohort.

### Data availability

The sncRNA-seq data that support the findings of this study are available from PPMI through the LONI data archive, https://www.ppmi-info.org/data. The authors declare that all the other data supporting the findings of this study are freely available.

### Code availability

The computer code written for the primary analysis is available upon request.

### Ethical compliance

The compliance with all ethical regulations is affirmed. Written and informed consent from each study participant of PPMI and NCER-PD has been collected prior to actions and analysis.

---

**Extended Data Figure 1.**
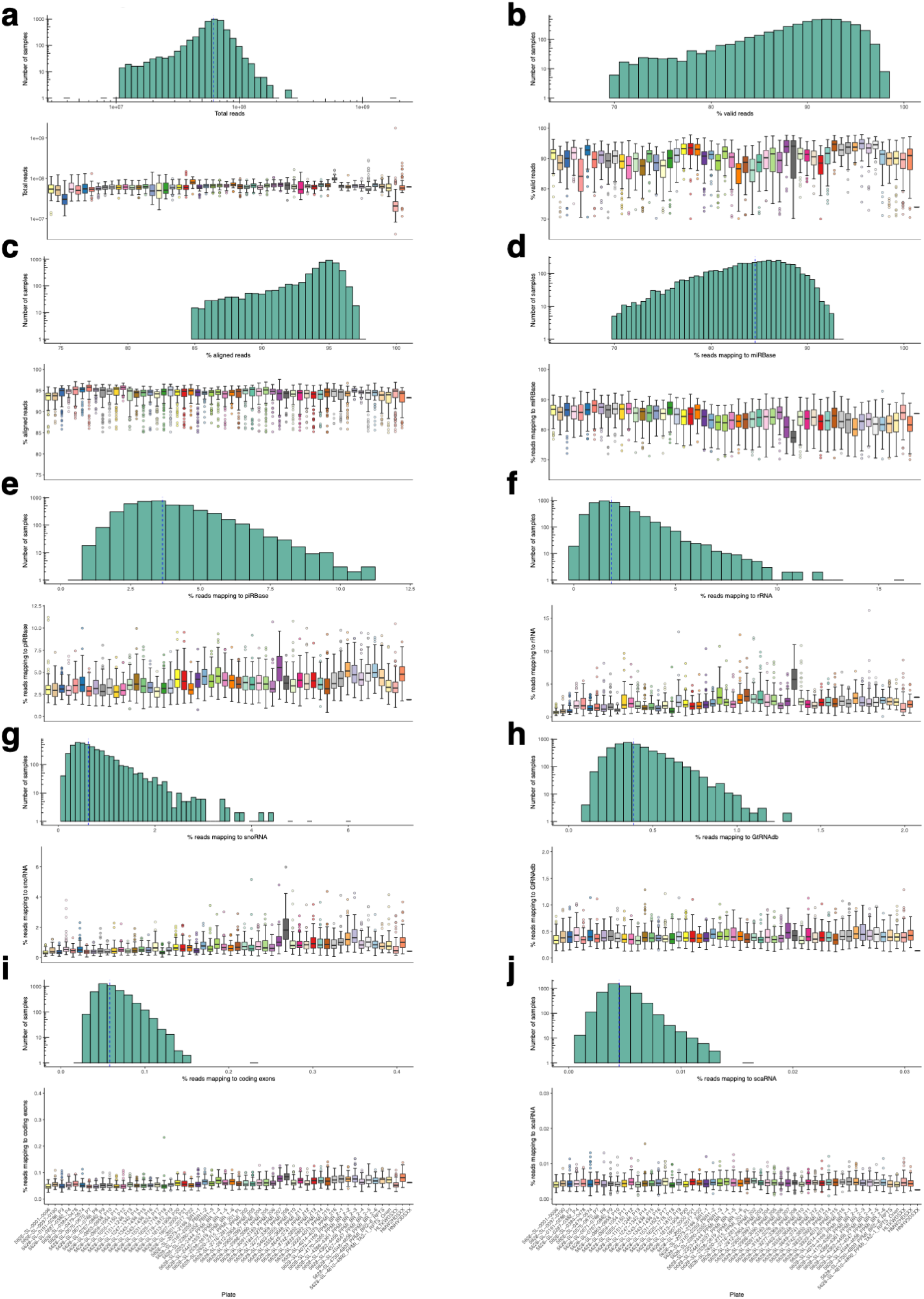
Detailed sncRNA-seq read statistics. **a-j)** Histogram plot for the number/percentage of reads (x-axis) and the number of samples (y-axis) on top, and boxplot for the distribution of reads per sequencing plate (x-axis) on the bottom. Each panel shows a different subset of reads using all valid samples. **a)** Shown are the total read counts. **b)** Shown are the percentage of valid reads. **c)** Shown are the percentage of reads aligned to the human genome. **d)** Shown are the percentage of reads mapping to miRBase human miRNA entries. **e)** Shown are the percentage of reads mapping to piRNA entries from piRBase. **f)** Shown are the percentage of reads mapping to ribosomal RNA. **g)** Shown are the percentage of reads mapping to small-nucleolar RNAs. **h)** Shown are the percentage of reads mapping to tRNA entries from GtRNAdb. **i)** Shown are the percentage of reads mapping to coding exons. **j)** Shown are the percentage of reads mapping to small cajal body-specific RNAs.

**Extended Data Figure 2.**
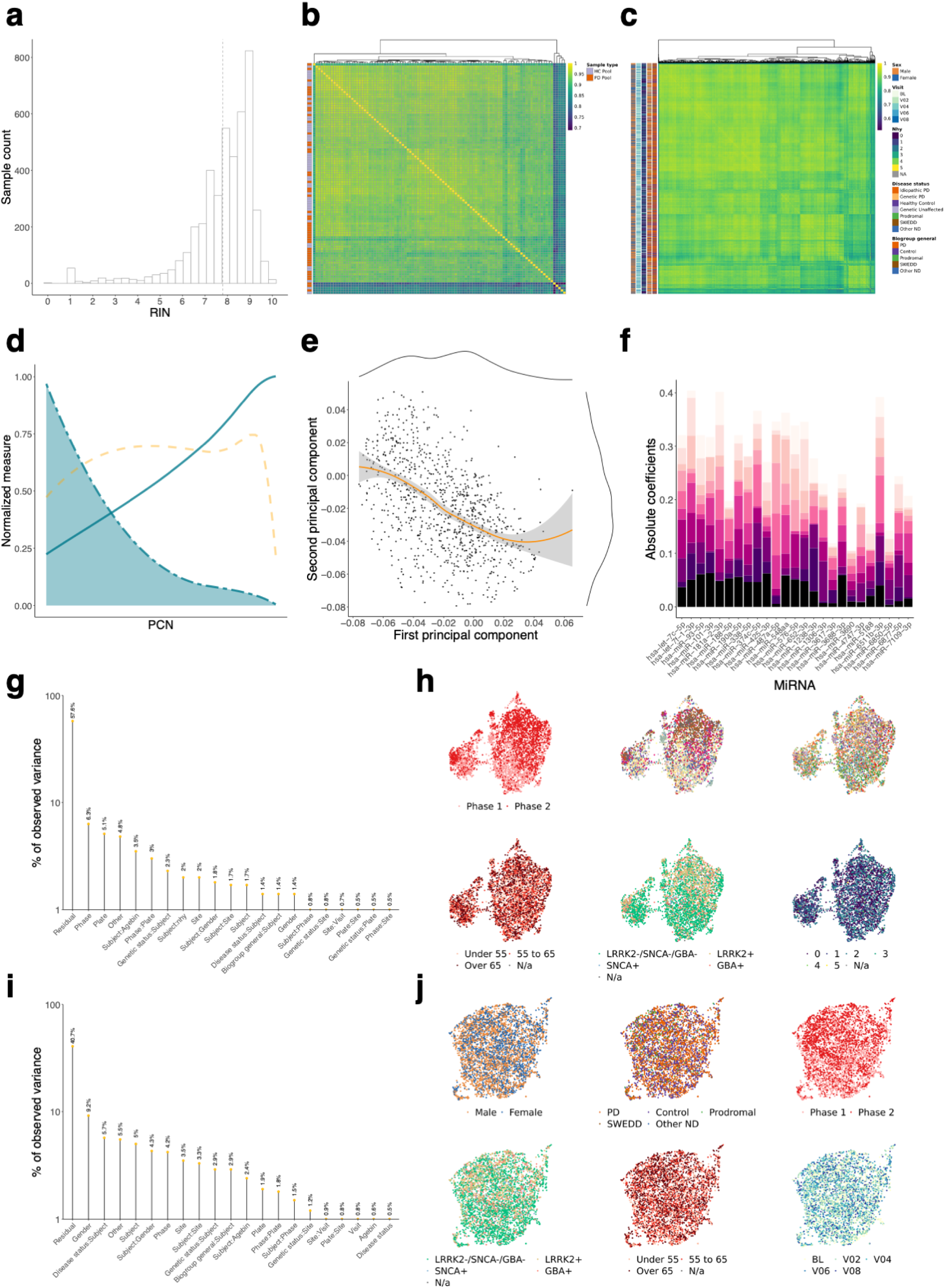
Quality control, batch variable analysis, and dimension reduction. **a)** Histogram of RNA integrity numbers (RINs) for all valid sequencing samples. **b)** Clustered pairwise correlation matrix of (pooled) technical controls and sequencing replicates based on the sncRNA counts. **c)** Clustered pairwise correlation matrix of all valid samples based on the known miRNA counts and with major annotation variables depicted on the left. **d)** The number of principal components resulting from PCA of the miRNA to sample expression matrix versus the cumulative percentage of variance explained (continuous blue line), the fraction of non-zero coefficients (dashed blue line), and the standard error (dashed orange line). **e)** Distribution of miRNA coefficient loadings onto the first two principal components. The smoothed trendline (orange) is enclosed by light grey bands showing the 95% confidence intervals. **f)** MiRNA versus stacked coefficient barplots for the miRNAs showing the highest sum of contributions along the top 10 principal components. **g)** PVCA based on sncRNA counts using the major annotation variables for all valid samples and combinations of such. **h)** UMAP embeddings for all valid samples using the sncRNA counts and colored in order by PPMI *project phase, sequencing plate, study participant, age binning, genetic status*, and *Hoehn and Yahr staging*. **i)** PVCA based on miRNA counts using the major annotation variables and combinations of such. **j)** UMAP embeddings for all valid samples using the miRNA counts and colored in order by *gender, biogroup*, PPMI *project phase, genetic status, age binning*, and *clinical visits*.

**Extended Data Figure 3.**
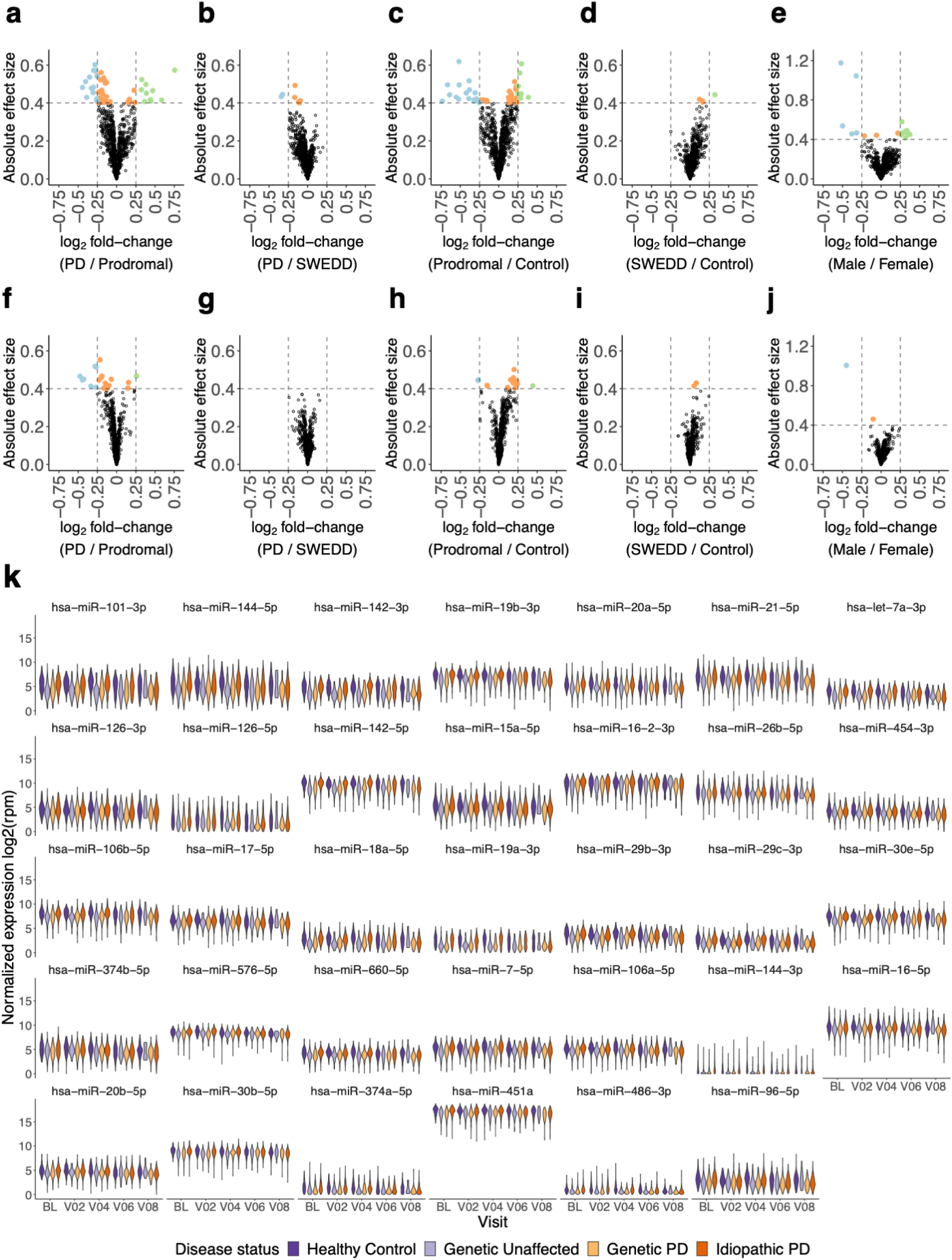
Complementary biogroup comparisons and miRNA expression distribution. **a-j)** Volcano plots with absolute effect size (cohen s d) of expressed miRNAs for the secondary group comparisons considered in the PPMI study. MiRNAs that exhibit both a considerable effect size and fold-change are colored in blue or green when being up- or down-regulated, respectively. Orange points depict miRNAs with a considerable effect size but a small fold-change. **k)** Normalized and log_2_-scaled expression of miRNAs showing a progressive depletion in PD (cf. Figure 3q)) wrapped by visit and disease status.

**Extended Data Figure 4.**
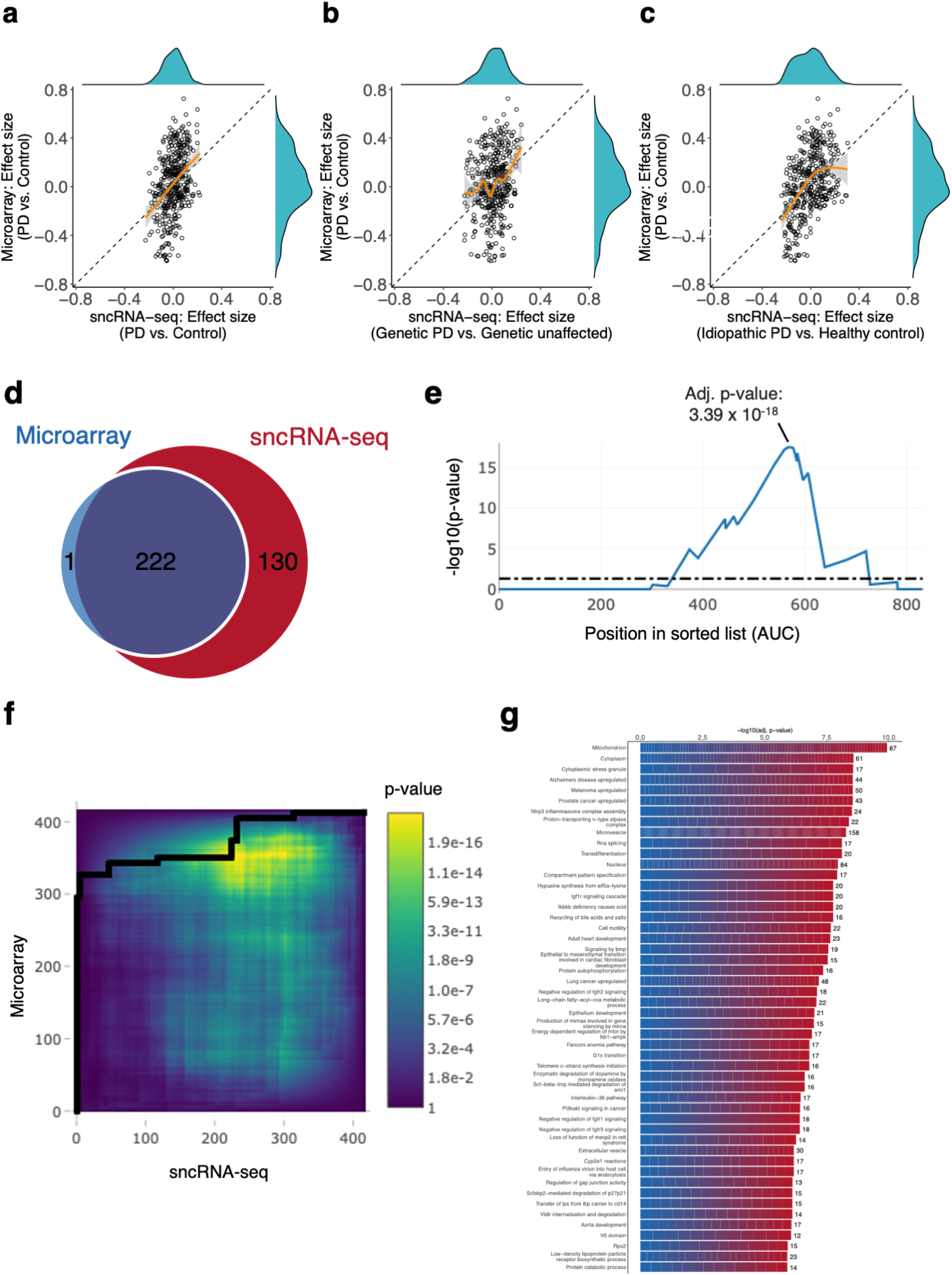
miRNA expression marker validation using 1, 440 microarray samples of the independent NCER-PD cohort. **a)** Scatterplot of miRNA effect size between total PD and controls obtained from sncRNA-seq and microarray. The orange trendline shows a generalized additive model (GAM) along with the 95% confidence intervals (grey shade). One-dimensional histograms are shown on the right and top for sncRNA-seq results and microarray results, respectively. **b)** Similar to **a)** but restricted to the effect sizes between genetic PD and unaffected genetic carriers from the sequencing study. **c)** Similar to **a)** but restricted to the effect sizes between idiopathic PD and healthy controls from the sequencing study. **d)** Venn-diagram for the most significant overlap between the miRNAs detected with sequencing or microarray as computed by DynaVenn. Input miRNAs were sorted by decreasing AUC for idiopathic PD vs. healthy controls in case of the sequencing study and PD vs. controls in case of the microarray study. **e)** The most significant overlap (adj. *p* = 3.39 *×* 10^−18^) was observed at *N* = 222. **f)** Two-dimensional representation of the entire p-value search space investigated by DynaVenn for all possible overlaps of miRNAs from the sequencing and microarray studies. **g)** Results of a miEAA 2.0 over-representation/enrichment analysis using the miRNAs determined for the most significant overlap displayed in **d)**. The x-axis shows the BH-FDR adjusted p-value for each category on the y-axis. The numbers on the right display the number of hits observed per category or biological pathway.

